# Neural activity and theta rhythmicity in the anterior hypothalamic nucleus are involved in regulating social investigation

**DOI:** 10.1101/2022.07.01.498407

**Authors:** Renad Jabarin, Wael Dagash, Shai Netser, Shelly Pal, Blesson K. Paul, Edi Barkai, Shlomo Wagner

## Abstract

Social interactions are highly complex, involving both approach and avoidance actions towards specific individuals, dependent on the social context. Currently, the brain regions subserving these behaviors are not fully known. The anterior hypothalamic nucleus (AHN) is a relatively unstudied and poorly defined brain area, known as part of the medial hypothalamic defensive system. Recent studies that examined the role of the AHN in various contexts have yielded contradicting results regarding its contribution to approach, avoidance, and escape behaviors. Yet, none of these studies has directly examined its role in social interactions. Here we explored the role of AHN neurons in regulating approach and avoidance actions towards distinct stimuli during various types of social interactions, using electrophysiological recording of neural activity in the AHN of behaving mice, c-Fos staining, and direct optogenetic stimulation. We found that theta rhythmicity in the AHN was enhanced during affiliative interactions, but decreased during aversive ones. Moreover, spiking activity of AHN neurons was found to be elevated more persistently during investigation of social stimuli, as compared to objects. Notably, AHN neuronal firing was found to be modulated by theta rhythmicity during social interactions. Finally, we found that during social interaction, direct optogenetic stimulation of AHN neurons augmented approach behavior towards stimuli associated with the optogenetic activation. Overall, our results suggest a context-dependent role for AHN neuronal activity in regulating approach behavior during social interactions, and for theta rhythmicity in mediating the valence of the social context.

## Background

Social interactions are highly complex, involving both approach and avoidance actions towards specific individuals. Thus, at a certain social context a subject may approach one individual (a friend) while avoiding a second one (a foe). Moreover, the same type of behavior may be driven by distinct intentions, such that a subject may approach an individual in either an affiliative or an offensive manner. These characteristics make the neurobiological mechanisms underlying social behavior difficult to elucidate (Adolphs, 2010). Nevertheless, during the last two decades, a plethora of studies have begun to reveal brain circuits subserving various types of social behavior (Ford and Young, 2021; Kohl and Dulac, 2018; McKinsey et al., 2018; Wei et al., 2021). These studies revealed the involvement of a vast network of mostly limbic brain regions, termed the social behavior network (SBN), in regulating mammalian social behavior (Dickinson et al., 2022; Goodson, 2005). Several of these regions reside in the hypothalamus, a brain structure known to regulate innate social behaviors, such as mating, aggression, and parenting (Anderson, 2012; Hashikawa et al., 2016; Kohl and Dulac, 2018; Matthews and Tye, 2019). However, it is likely that additional brain regions are involved in regulating social behavior.

One such brain region with a potential role in regulating social interactions is the anterior hypothalamic nucleus (AHN, also known as anterior hypothalamic area - AHA). The AHN is a relatively understudied and poorly defined brain area, known as part of the medial hypothalamic defensive system (Canteras, 2002). It includes mainly inhibitory GABAergic neurons (Bang et al., 2022; Wang et al., 2015; Xie et al., 2022) which largely project to several brain regions thought to be central to fight-or-flight behavior, such as the lateral septum (LS) and dorsal periaqueductal gray (dPAG) area (Risold et al., 1994; Xie et al., 2022). In rats and mice, the AHN was shown to take part in responses to predators (Cezario et al., 2008; Martinez et al., 2008; Mendes-Gomes et al., 2020) as well as in regulating parental behavior (Tachikawa et al., 2013). In recent years, several studies explored the role of AHN neurons in the context of anxiety and avoidance behaviors, using advanced techniques (e.g., optogenetics), with contradicting results (Anthony et al., 2014; Bang et al., 2022; Wang et al., 2015). Yet, none of these studies examined the role of the AHN in the context of social behavior. Another recent study (Xie et al. 2021) demonstrated that GABAergic neurons in the AHN are involved in attack behavior which follows physical noxious stimuli. Intriguingly, this study reported that in a social setting, optogenetic activation of GABAergic AHN neurons reduces aggression and enhances social investigation, thus suggesting a distinct role for the AHN in social contexts. These results suggest that the role of the AHN markedly differs in distinct contexts.

Here we hypothesize that the behavioral contribution of the AHN is context-dependent and that it’s role differ in social context than in other contexts such as during predator’s attack. To explore the behavioral contribution of the AHN in social context, we explored the role of AHN neurons in regulating approach and avoidance actions towards distinct stimuli during social discrimination tasks. To that end, we used electrophysiological recording of neuronal population activity in the AHN, c-Fos staining, and direct optogenetic stimulation. We found that theta rhythmicity in the AHN was enhanced during affiliative, but decreased during aversive social interactions and that spiking activity of AHN neurons was elevated more persistently during investigation of social stimuli, as compared to objects. Notably, spiking activity of AHN neurons was found to be modulated by LFP theta rhythmicity during social interactions. Finally, we found that during social interaction, direct optogenetic stimulation of AHN neurons augmented approach behavior towards stimuli associated with this stimulation. Overall, our results suggest a role for AHN neuronal activity in regulating approach behavior during social interactions, and for theta rhythmicity in mediating the valence of the social context. They also propose that the AHN role is highly context-dependent.

## Results

Theta and gamma power of local field potential in the AHN increase during social interaction

For recording neural activity from the AHN of behaving mice (n=17), we chronically implanted tetrodes (4x4; Fig. S1A) in this brain area and conducted behavioral experiments, while recording electrophysiological signals. Each experiment comprised a 5-min social preference (SP) session (Fig. 1A, B), followed 30 min later by a 5-min session of free social interactions (Fig. S2 A-B). For each electrophysiological recording we separately analyzed local field potential signals (LFP; low-pass filtered at 0.3 kHz) and spiking activity (band-pass filtered at 0.3-5 kHz). LFP signals were analyzed first by generating a power spectral density (PSD) profile separately for each of the three 5-min stages of the session: Pre-encounter, Encounter, and Post-encounter. As apparent in the single-session example displayed in Fig. 1C, while the Pre-encounter and Post-encounter profiles were almost identical, the Encounter profile reflected increased power in both the theta (4-10 Hz) and gamma (30-80 Hz) bands, suggesting enhanced theta and gamma rhythmicity in the AHN during social interaction.

**Figure 1.**
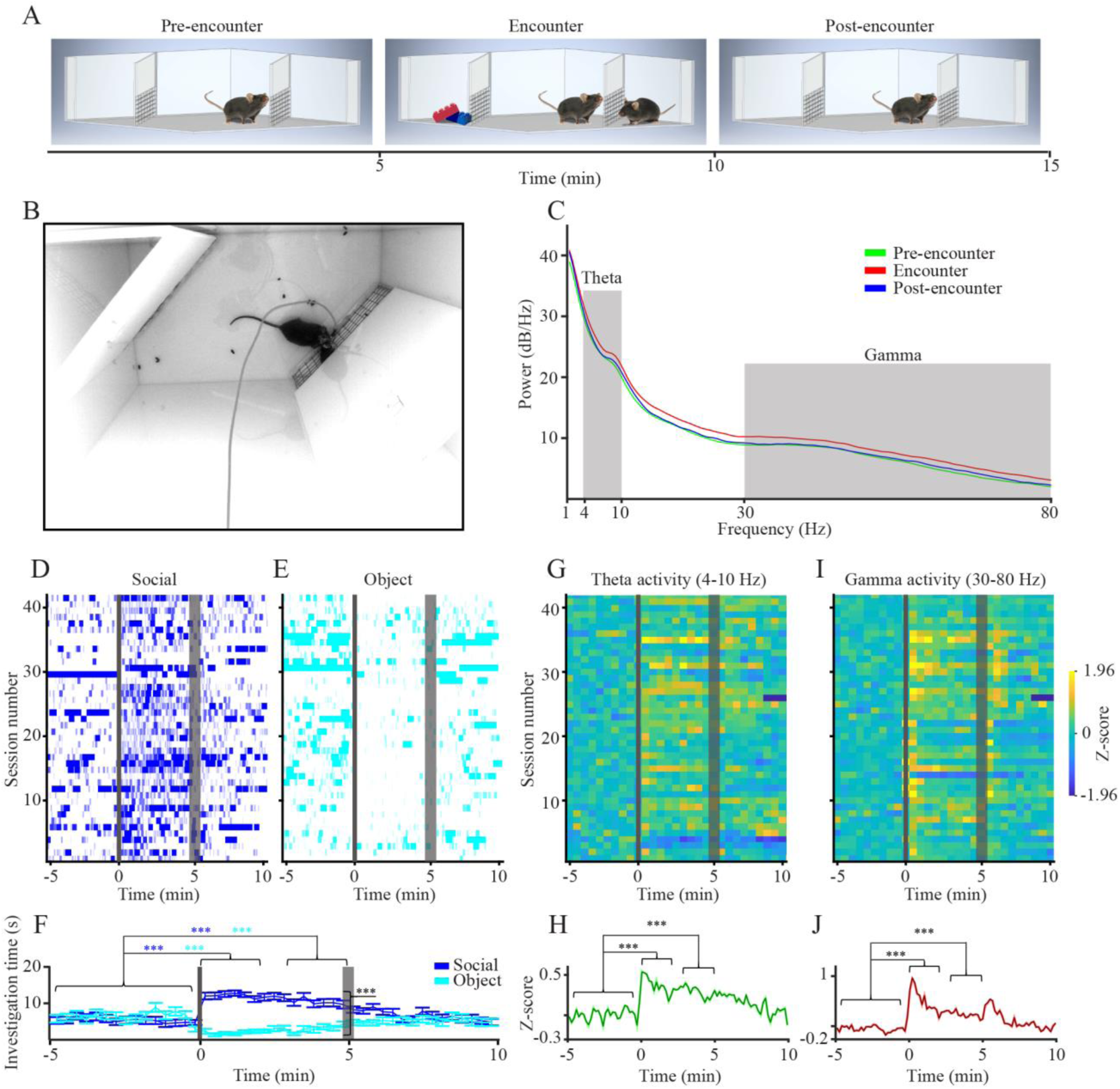
AHN theta and gamma rhythms are enhanced during the social preference (SP) test. (A) Schematic depiction of the three stages of the test. (B) A picture of the experimental arena during the SP test. (C) Power spectral density (PSD) profiles of local field potential (LFP) signals recorded during each of the three 5-min stages of the test from a single animal, color-coded according to the test stage. The theta (4-10 Hz) and Gamma (30-80 Hz) ranges are labeled in gray. (D) Behavioral raster-plots of investigation bouts towards social stimuli along 42 sessions. Gray bars represent the time of insertion (thin bar) and removal (thick bar) of the stimuli from the chambers. Time 0 represents the beginning of the test. (E) As in D, for object stimuli during the same sessions. (F) Mean investigation time (±SEM, 20-s bins) during the same sessions shown in D-E, plotted separately for the social (blue) and object (light blue) stimuli. Note the statistically significant change from the 5-min Baseline, in investigation time of both social (increase) and object (decrease) stimuli during the first and last two minutes of the Encounter. (G) Heat-map of Z-score analysis of theta power of LFP signals recorded during the same sessions shown in D-E. (H) Mean (±SEM) Z-score analysis of theta power of LFP signals recorded during the same sessions shown in D-E. (I) As in F, for gamma power. (J) As in H, for Z-score of gamma power. ***p<0.001, paired t-test with Bonferroni correction for multiple comparisons following main effect of time in ANOVA. See also Fig. S2.

**Figure 2.**
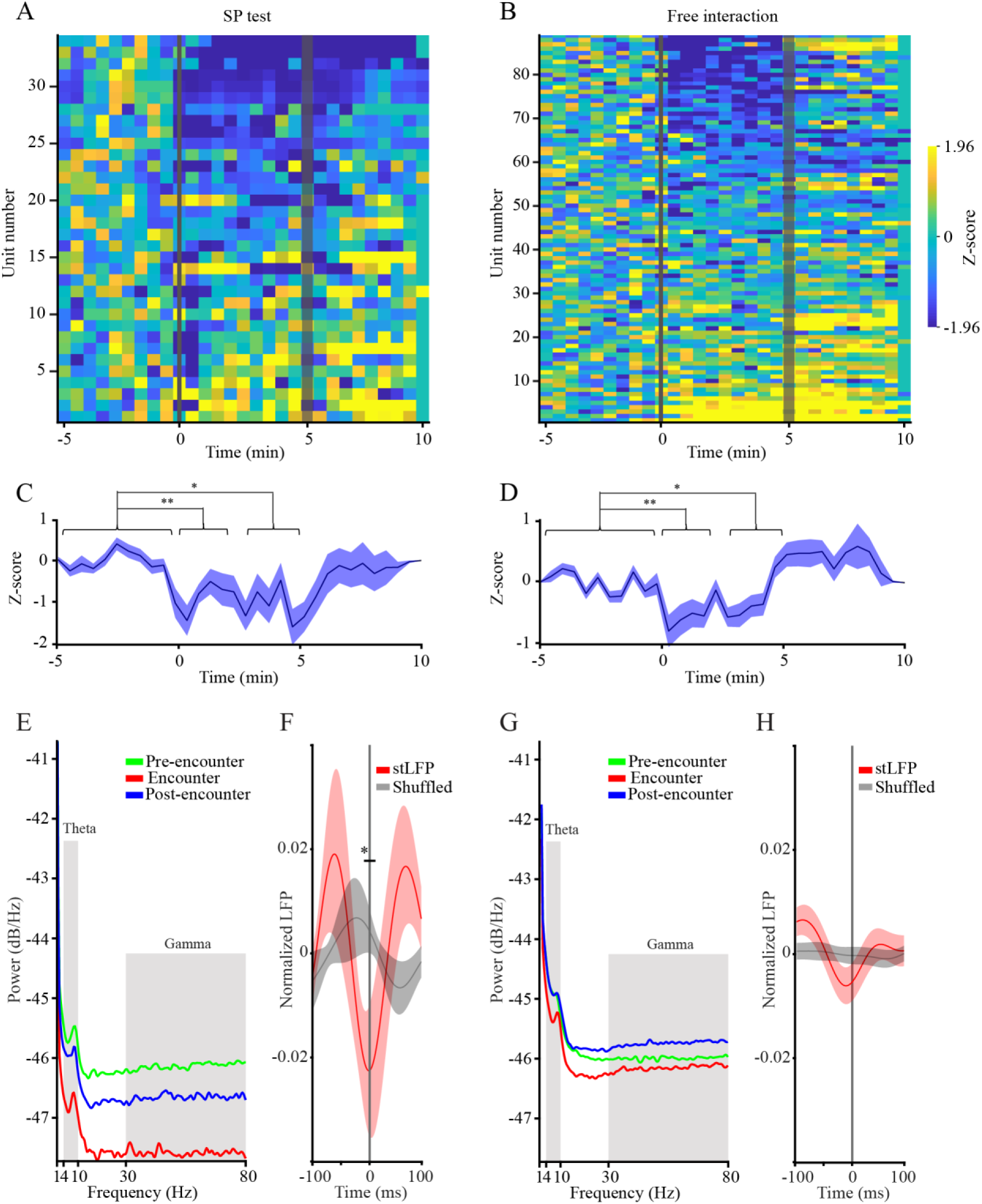
AHN Spiking activity is generally reduced and synchronizes with the LFP theta rhythm during social encounter. (A) Heat-maps of Z-score analysis of multi-unit activity (MUA) for all units recorded during SP tests, sorted from inhibition (top) to excitation (bottom). (B) As in A, for free interaction sessions. (C) Mean (±SEM) Z-score analysis of the units shown in A. Note the statistically significant decrease in mean spiking activity during the first (left) and last (right) two minutes of the Encounter, as compared to the 5-min Baseline stage. *p<0.05, **p<0.01, Wilcoxon test for non-parametric paired comparisons following main effect in Friedman’s test. (D) As in C, for the units shown in B. (E) PSD profiles of the MUA recorded during each stage of the SP test, color-coded according to the test stage. The theta and gamma ranges are labeled in gray. Note the general reduction during the Encounter stage, which is caused by the lower mean firing rate at this stage, as compared to the other stages. Also, note the clear peak at the theta range, demonstrating theta rhythmicity of spiking activity. (F) Spike-triggered LFP (stLFP, ±SEM) of theta-filtered LFP signals (red) and of shuffled LFP signals (gray) for the Encounter period. Note the statistically significant difference between the mean stLFP and shuffled signals at time 0 (between -10 to 10 ms), suggesting synchronization of spiking activity and LFP theta rhythmicity. (G) As in E, for free-interaction experiments. (H) As in F, for free-interaction experiments. Note that here the difference between STA and shuffled signals is not statistically significant, but keeps the same trend as for the SP sessions shown in F. See also Fig. S1.

We analyzed the behavior of the subjects during the SP task using our published automated analysis system (Netser et al., 2019). This system allows precise tracking of investigation bouts conducted by the subject mouse towards each of two chambers containing either a novel conspecific (social stimulus) or an object (Netser et al., 2017). As apparent in Fig. 1 D-F, while the subjects did not prefer any of the empty chambers in the Pre-encounter stage, they showed a clear preference for investigating the chamber of the social stimulus during the Encounter stage (Wilcoxon’s test: Z=-5.345, p<0.001). At the Post-encounter stage, for a while, the subjects kept a preference for the empty chamber that previously contained the social stimulus. This preference gradually disappeared [2-way repeated measures ANOVA-Time: F(1.64,67.4)=8.084, p<0.01; Stimulus: F(1,41)=63.763, p<0.001; Time x Stimulus: F(1.68,69.164)=49.484, p<0.001; *post hoc* paired t-test with Bonferroni correction for multiple comparisons - Social (pre-encounter-first 2 min): t(41)=-9.319, p<0.001; Social (pre-encounter-last 2 min): t(41)=-6.758, p<0.001; Object (pre-encounter-first 2 min): t(41)=6.178, p<0.001; Object (pre-encounter-last 2 min): t(41)=4.673, p<0.001; First 2 min (Social-Object): t(41)=13.672, p<0.001; Last 2 min (Social-Object): t(41)=6.945, p<0.001].

We used Z-score analysis to normalize the theta and gamma power of the LFP signals recorded during the various experiments, with the Pre-encounter stage serving as a baseline (Fig. 1G-J). We found a clear increase in both theta and gamma power following the introduction of the social stimulus to the arena, compared to the Pre-encounter stage (Fig. 1G-J). This increase was statistically significant for both the first two minutes and last two minutes of the session [Theta: Repeated Measures ANOVA-F(2,82)=19.1810, p<0.001; Bonferroni *post hoc* comparisons-Pre-encounter-first 2 min: p<0.001; Pre-encounter-last 2 min: p<0.01; First 2 min-Last 2 min: p=0.095; Gamma: Friedman’s test - *χ*^2^=53.476, p<0.001; Wilcoxon test- Pre-encounter-first 2 min: Z= -0.64, p<0.001; Pre-encounter-last 2 min: Z=-6.283, p<0.01; First 2 min-Last 2 min: Z=-3.047, p<0.01]. Similar results were observed for the same parameters recorded from the same animals during free social interactions [Fig. S2E-H; Theta: Repeated Measures ANOVA-F(2,116)=53.684; p<0.001; Bonferroni *post hoc* comparisons: Pre-encounter-first 2 min: p<0.001; Pre-encounter-last 2 min: p<0.01; First 2 min-Last 2 min: p=0.543; Gamma: Friedman’s test - *χ*^2^=75.76, p<0.001, Wilcoxon test: Pre-encounter-first 2 min: Z= -5.645, p<0.001; Pre-encounter-last 2 min: Z=-4.33, p<0.01; First 2 min-Last 2 min: Z=-4.62, p<0.001].

### Spiking activity in the AHN is generally reduced but gets more rhythmic during social interactions

For analyzing the spiking activity recorded during the SP and free interaction experiments (shown in Fig. 1 and Fig. S2, respectively), this activity was sorted into single- and multi-units (SUA and MUA, respectively; Fig. S1B, D). Notably, since we managed to reliably separate a relatively low number of SUAs (2 out of 34 for SP and 33 out of 89 for free interaction), we pooled all SUAs and MUAs together and considered them all as MUAs. As apparent in Fig. S1E, while analyzing the firing rate across each session, we observed variable responses of these MUAs. For example, some MUAs responded to the social stimulus insertion with increased activity (Fig. S1E, 1^st^ trace), some responded with transient inhibition (2^nd^ trace), some increased their activity only after stimulus removal (3^rd^ trace) and some did not respond at all (4^th^ trace). This variability was also demonstrated by the heat-maps representing the Z-score analysis of all MUAs recorded during SP (Fig. 2A) and free interaction (Fig. 2B) sessions. However, when the mean Z-score of all MUAs was calculated, we observed a general inhibition in firing rate for both SP (Fig. 2C; Friedman’s test: *χ*^2^=11.146, p<0.01) and free interaction (Fig. 2D; Friedman’s test: *χ*^2^=5.647, p<0.01) sessions. This inhibition was statistically significant, compared to baseline, for both the first two minutes and last two minutes of both types of session [SP: Wilcoxon’s test: Pre-encounter-first 2 min: Z= -3.206, p<0.01; Pre-encounter-last 2 min: Z=-2.48, p<0.05; First 2 min-Last 2 min: Z=-1, p=0.951; Free Interaction: Wilcoxon’s test: Pre-encounter-first 2 min: Z=-3.256, p<0.01; Pre-encounter-last 2 min: Z= -2.69, p<0.05; First 2 min-Last 2 min: Z=-1.782, p=0.225]. Thus, social interaction is accompanied by a general reduction in the firing rate of AHN neurons.

Despite the general reduction in spiking activity, it may be possible that during social interactions, when the power of LFP theta and gamma rhythmicity is high, spiking activity is synchronized by these rhythms to create more coherent firing of the neuronal population (Fries, 2015). To examine this possibility, we first calculated the PSD profiles of the spiking activity for the various stages of the SP paradigm. While the general power was reduced during the encounter due to the general reduction in firing, a clear peak at the theta range was apparent, suggesting significant theta rhythmicity of spiking activity (Fig. 2E). To examine possible synchronization between LFP theta rhythmicity and firing activity, we conducted spike-triggered LFP averaging (stLFP) of the theta-filtered LFP signals and compared it to shuffled signals. We found that during the Encounter period there was a significant difference between the stLFP and the shuffled LFP profile, suggesting preferred firing at the trough of the LFP theta rhythm (Fig. 2F, Paired samples t-test: t(26)=2.1557, p<0.05). Qualitatively similar results were obtained during free interaction sessions, although in this case the difference between the stLFP and the shuffled LFP profile was not statistically significant (Fig. 2G-H, Paired samples t-test: t(86)=1.4427, p=0.15). Overall, these results suggest synchronization between the firing activity and the LFP theta rhythmicity during the social encounter.

### AHN spiking activity correlates with stimulus investigation in a stimulus-dependent manner

Next, we examined each of the recorded parameters (theta power, gamma power, and firing rate) in correlation with the specific investigation bouts, analyzed separately for each stimulus (object and social). Such an analysis can be done reliably only for SP sessions, while in free interaction sessions it is difficult to determine the exact starting point of each investigation bout. As for both theta and gamma power, while there seems to be a weak increase in the Z-score in response to the encounter with the social and object stimuli, respectively, during the first two seconds following the bout beginning, this increase was hardly significant and no significant difference was observed between bouts towards social and object stimuli [Fig. 3A-F, Theta: two-way Repeated Measures ANOVA: Stimulus: F(2,80)=3.787, p<0.05; Time: F(1,40)=1.323, p=0.257; Stimulus x Time: F(2,80)=0.567, p=0.567; *post hoc* paired t-test with Bonferroni correction for multiple comparisons: Social (Baseline-First 2 s): t(41)=-2.455, p=0.108; Social (Baseline-Last 2 s): t(41)=0.705, p=1; Object (Baseline-First 2 s): t(40)=-1.453, p=0.924; Object (Baseline-Last 2 s): t(40)=0.799, p=1; First 2 s (Social-object): t(40)=0.954, p=1; Last 2 s (Social-Object): t(40)=0.959, p=1; Gamma: two-way Repeated Measures ANOVA: Stimulus: F(1.7,68.9)=8.72, p<0.05; Time: F(1,40)=0.002, p=0.968; Stimulus x Time: F(2,80)=0.493, p=0.613; *post hoc* paired t-test with Bonferroni correction for multiple comparisons: Social (Baseline-First 2 s): t(41)=-2.489, p=0.102; Social (Baseline-Last 2 s): t(41)=-0.886, p=1; Object (Baseline-First 2 s): t(40)=-2.938, p<0.05; Object (Baseline-Last 2 s): t(40)=-0.016, p=1; First 2 s (Social-object): t(40)=-0.574, p=1; Last 2 s (Social-Object): t(40)=0.488, p=1]. In contrast, when MUA firing rate was analyzed using the same methodology, we found a statistically significant change, compared to baseline, for investigation bouts towards both the social and object stimuli [Fig. 3G-I; Friedman’s test-Stimulus: *χ*^2^=5.647, p<0.05; Time: *χ*^2^=19.993, p<0.001; Wilcoxon’s test with Bonferroni correction for multiple comparisons - Social (Baseline-First 2 sec): z=-3.24, p<0.01; Social (Baseline-Last 2 sec): z=-0.983, p=1; Social (First 2 s-Last 2 s): z=-1.714, p=0.56; Object (Baseline-First 2 s): z=-2.248, p=0.2; Object (Baseline-Last 2 s): z=-2.6, p=0.08; Object (First 2 s-Last 2 s): z=-4.026, p<0.001]. Yet, while social investigation bouts were characterized by a significant increase in firing rate, investigation of object stimuli elicited an initial increase in Z-score, followed by a significant decline 4-6 s after the beginning of the investigation bout. Consequently, a statistically significant difference between responses to social and to object stimuli was observed 4-6 s after the beginning of the investigation bout [Fig. 3I; Wilcoxon’s test with Bonferroni correction for multiple comparisons: First 2 s: z=-0.82, p=1; Last 2 s: z=-2.898, p<0.05]. Altogether, these results suggest that while theta or gamma rhythms in the AHN are related to the social context, and therefore show a general increase during social interaction, spiking activity correlates with stimulus investigation, in a stimulus-dependent manner.

**Figure 3.**
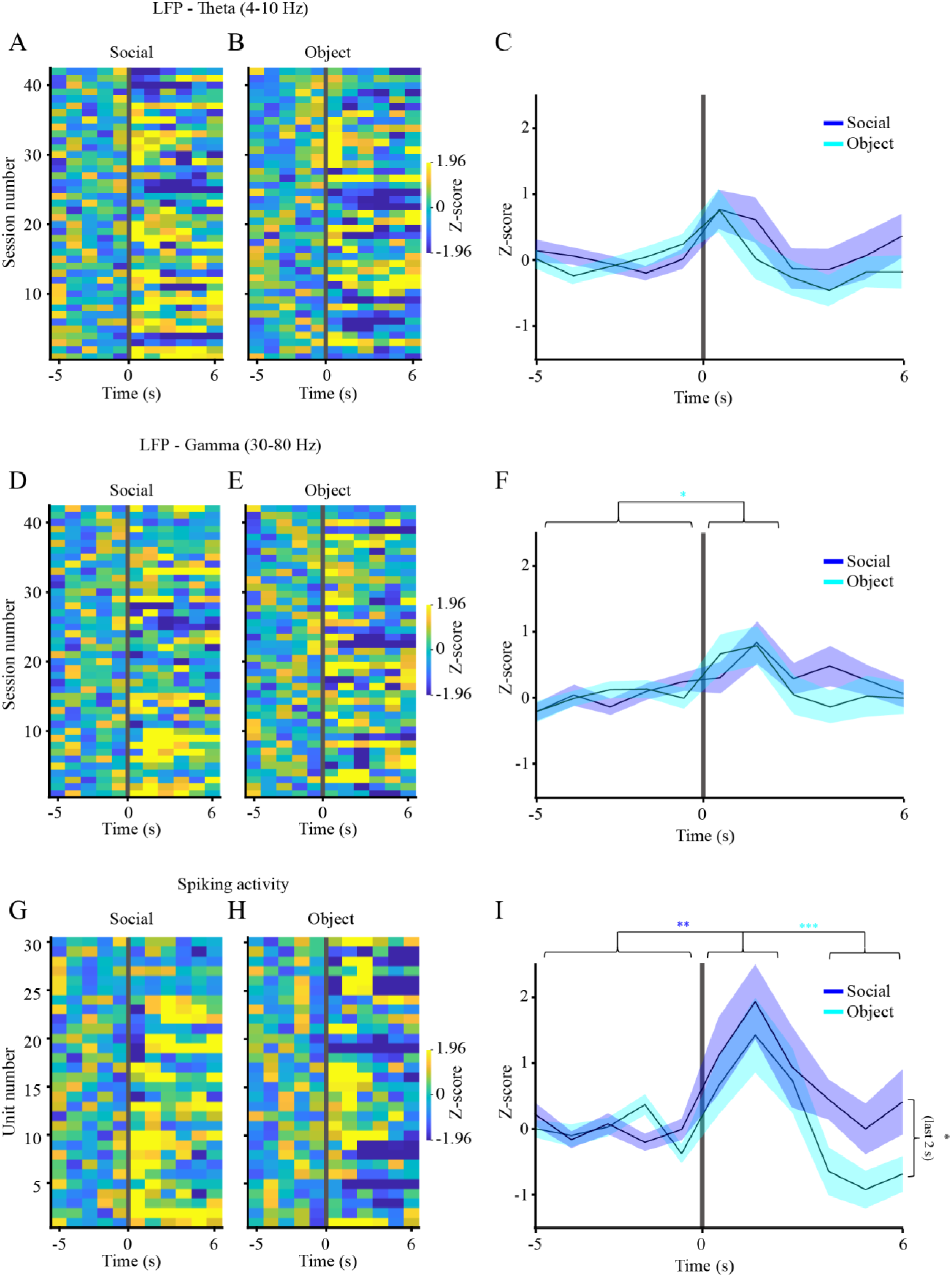
Spiking activity, but not LFP rhythms, is specifically enhanced during social investigation bouts. (A) Heat maps of mean Z-score of LFP theta power recorder from 5-s before to 6-s after the beginning of social investigation during SP sessions. Each line represents the mean value averaged across all bouts for a single session. (B) As in A, for object investigation bouts during the same sessions. (C) Mean (±SEM) Z-score analysis for all the sessions shown in A-B, averaged separately for social (blue) and object (light blue) stimuli. No significant change was observed, as compared to the baseline (5 s before beginning of bout) period. (D) As in A, for gamma power. (E) As in B, for gamma power. (F) As in C, for gamma power. Note that in this case a slightly significant increase was observed for the first two seconds of the bout towards the object stimulus, as compared to the baseline. (G) As in A, for MUA. (H) As in B, for MUA. (I) As in C, for MUA. Note that a significant change from baseline was found for social investigation during the first two seconds. Note also that a significant difference was observed between social and object investigation bouts during the last two seconds. *p<0.05, **p<0.01, ***p<0.001, Wilcoxon test for non-parametric paired comparisons with Bonferroni correction for multiple comparisons following main effect in Friedman’s test.

### AHN theta rhythmicity of LFP signals differs between attractive and aversive social stimuli

So far, we have explored AHN activity during interactions of the subject animals with stimuli which are both attractive, although at different levels of attractiveness (social vs. object stimuli). Yet, avoidance behavior is usually exhibited towards aversive stimuli. Thus, it may be that neural activity in the AHN differs between attractive and aversive stimuli. To examine this possibility in a social context, we employed a novel behavioral paradigm that enables modifying the valence of social stimuli. In this stimulus-specific social fear conditioning (SFC) paradigm (schematically described in Fig. 4A and in more detail in Fig. S3A), each subject (n=7) performed two consecutive SP tests (Baseline tests, separated by 15 min) before the SFC session. For each of these tests, we used a social stimulus of a different mouse strain (C57BL/6J and ICR), so as to enhance the ability of a subject to discriminate between them. Twenty minutes after the second SP test, we conducted a 5-min SFC session using the same ICR stimulus used for the Baseline test, but in a different spatial context. Twenty minutes later, we conducted two SP tests with the same social stimuli as used before the SFC session (Early recall tests). The same tests with the same stimuli were then repeated 24 h later (Late recall tests). Examples of subject’s movement tracking during each of the paradigm stages are shown in Fig. 4B. To examine whether social-preference was significantly modified by the social fear conditioning, we conducted a 2 strains (C57BL/6J vs. ICR) x 2 stimuli (Empty vs. Social) two-way mixed model ANOVA, for each of the stages. As for the Baseline stage, we found a significant main effect only for the stimulus, and a similarly significant preference was exhibited by the subjects towards both C57BL/6J and ICR stimuli, as compared to objects [Fig. 4C, two-way mixed model ANOVA: Stimulus: F(1,10)=78.494; p<0.001; Strain: F(1,10)=2.522, p=0.143, Stimulus x Strain: F(1,10)=3.121, p=0.108; *post hoc* paired t-test with Bonferroni correction for multiple comparisons: ICR: t(6)=4.423, p<0.01; C57BL/6J: t(4)=16.174, p<0.001]. In contrast, during Early recall sessions, we found a significant interaction between stimulus and strain: while the subjects did not show any preference for C57BL/6J stimuli over objects, they developed a marginally significant avoidance towards the fear-conditioned ICR stimulus [Fig. 4D, two-way mixed model ANOVA-Stimulus: F(1,11)=1.293; p=0.28; Strain: F(1,11)=0.015, p=0.906, Stimulus x Strain: F(1,11)=6.645, p<0.05; *post hoc* paired t-test with Bonferroni correction for multiple comparisons: ICR: t(5)=-2.858, p=0.07; C57BL/6J: t(6)=0.976, p=0.734]. Moreover, at the Late recall stage, we found again a significant interaction between stimulus and strain, with *post hoc* analysis revealing a social preference for the C57BL/6J only [Fig. 4E, two-way mixed model ANOVA: Stimulus: F(1,7)=1.565; p=0.25; Strain: F(1,11)=6.579, p<0.05, Stimulus x Strain: F(1,7)=9.092, p<0.05; *post hoc* paired t-test with Bonferroni correction for multiple comparisons: ICR: t(3)=-0.983, p=1; C57BL/6J: t(4)=3.934, p<0.05]. Thus, while in the Baseline tests the subjects showed similar preference for both social stimuli, during the Late recall experiments, their behavior depended on the strain of the social stimulus.

**Figure 4.**
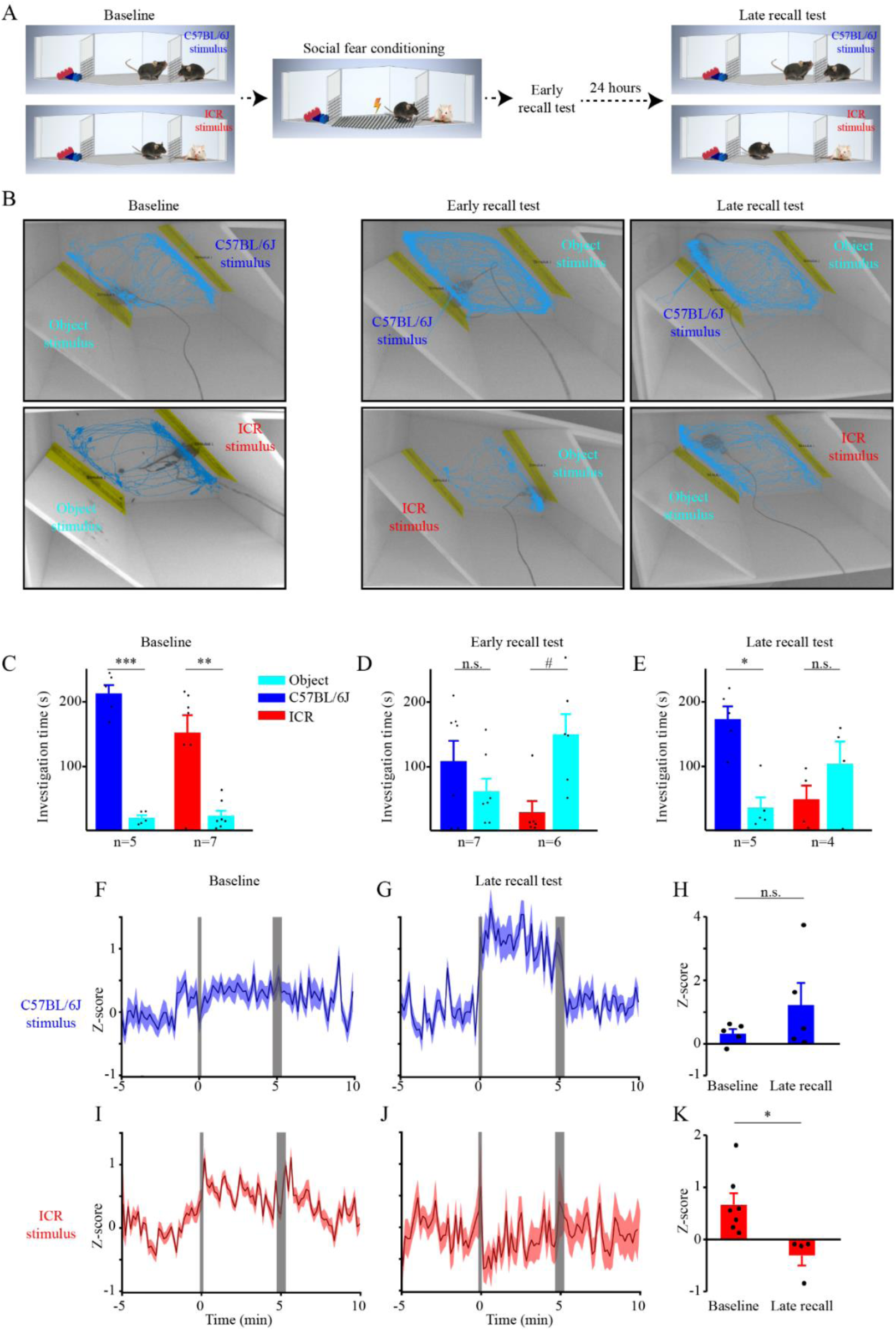
Theta power of LFP signals in the AHN increases during affiliative social interactions and decreases during aversive ones. (A) A schematic depiction of the social fear conditioning (SFC) paradigm. (B) Representative images of the arena and tracked movement of subject animals during the various stages of the SFC paradigm. The three stimuli are color coded (C57BL/6J – blue, ICR – red, object – light blue). (C) Mean (±SEM) Investigation time dedicated to each of the stimuli during the Baseline SP tests. Note the similar results obtained with both C57BL/6J and ICR stimuli. (D) As in C, for Early recall tests. (E) As in C, for Late recall tests. Note the normal preference for the C57BL/6J mice, in contrast to the fear-conditioned ICR stimulus that draws no preference as compared to object stimuli. (F) Mean (±SEM) Z-score analysis of theta power along the time course of the Baseline SP test using C57BL/6J mice. (G) As in F, for the Late recall test using the same stimulus. (H) Mean Z-score analysis of the change in theta power during the Encounter stage of the SP test using C57BL/6J stimuli, calculated separately for Baseline (left) and Late recall (right) experiments. Note the apparent, but not statistically significant increase in theta power during Late recall, as compared to Baseline. (I) As in F, for ICR stimuli. (J) As in G, for ICR stimuli. (K) As in H, for ICR stimuli. Note the statistically significant reduction in theta power during the Late recall test, as compared to Baseline. #=0.07,*p<0.05, **p<0.01, ***p<0.001 t-test with Bonferroni correction for multiple comparisons following a significant effect in ANOVA. See also Fig. S3.

When analyzing the theta rhythmicity recorded in the AHN during the Baseline and Late recall tests (using Z-score analysis), we found a significant interaction between stage and strain [two-way mixed model ANOVA: Strain: F(1,7)=4.691; p=0.069; Stage: F(1,7)=0.264, p=0.623, Stimulus x Strain: F(1,7)=6.223, p<0.05]. While during Baseline, theta rhythmicity trended to be similarly elevated during sessions with both C57BL/6J and ICR stimuli (Fig. 4F-H, independent samples t-test: C57BL/6J: t(4.34)=-1.266, p=0.269), this was not the case during Late recall sessions. At this stage, we observed an apparently augmented theta rhythmicity towards the C57BL/6J stimulus, as compared to Baseline, as opposed to a significantly reduced rhythmicity towards the fear-conditioned ICR stimulus (Fig. 4I-K; independent samples t-test: t(9)=2.962, p<0.05). In contrast, no significant difference was observed between strains or stimuli for gamma rhythmicity [Fig. S3B-G, two-way mixed model ANOVA: Strain: F(1,7)=0.327; p=0.585; Stage: F(1,7)=0.05, p=0.83, Stimulus x Strain: F(1,7)=2.192, p=0.182].

Thus, theta, but not gamma rhythmicity in the AHN seems to reflect the valence of the social stimulus during a social encounter, by showing enhancement during interaction with attractive stimuli, as compared to reduction in the presence of aversive stimuli.

### AHN c-Fos expression following social encounter does not differ between attractive and aversive stimuli

Unfortunately, we did not have enough MUA recorded during these experiments in order to reliably assess the spiking activity. Therefore, to examine the possibility that AHN neural activity during Late recall sessions differs between fear-conditioned and neutral social stimuli, we used c-Fos immunostaining as a proxy for neural activity. These experiments (n=14) were conducted with BALB/cJ mice serving as fear-conditioned stimuli instead of ICR mice. This strain change did not alter the behavioral results [Fig. 5A-C, *Baseline:* two-way mixed model ANOVA: Stimulus: F(1,36)=23.352; p<0.001; Strain: F(1,36)=0.799, p=0.377, Stimulus x Strain: F(1,36)=0.751, p=0.392; *post hoc* paired t-test with Bonferroni correction for multiple comparisons: BALB/c: t(18)=2.693, p<0.05; C57BL/6J: t(18)=4.212, p<0.01; *Early Recall-*two-way mixed model ANOVA: Stimulus: F(1,26)=5.805; p<0.05; Strain: F(1,26)=1.173, p=0.289, Stimulus x Strain: F(1,26)=4.85, p<0.05; *post hoc* paired t-test with Bonferroni correction for multiple comparisons: BALB/c: t(13)=-3.285, p<0.05; C57BL/6J: t(13)=-0.146, p=1; *Late Recall:* Stimulus: F(1,12)=0.000; p=0.984; Strain: F(1,12)=1.215, p=0.292, Stimulus x Strain: F(1,12)=12.984, p<0.01; *post hoc* paired t-test with Bonferroni correction for multiple comparisons: BALB/c: t(7)=-2.19, p=0.13; C57BL/6J: t(5)=5.186, p<0.01], thus supporting the validity of the SFC paradigm across distinct strains of stimuli. When analyzing c-Fos expression in the AHN following a single Late recall test, we found a significantly higher number of c-Fos positive cells for both types of social stimuli, compared to control (no social stimulus during the Late recall test), with no significant difference between the fear-conditioned BALB/cJ and the neutral C57BL/6J stimuli (Fig. 5D-E, Kruskal Wallis test: H=21.53, p<0.001; Dunn’s multiple comparisons: Control vs. C57BL/6J: p<0.01; Control vs. BALB/c, p<0.001; C57BL/6J vs. BALB/c, p=0.308). These results suggest that it is the theta rhythmicity in the AHN, rather than the level of spiking activity, which reflects the valence of the social stimulus during social encounters.

**Figure 5.**
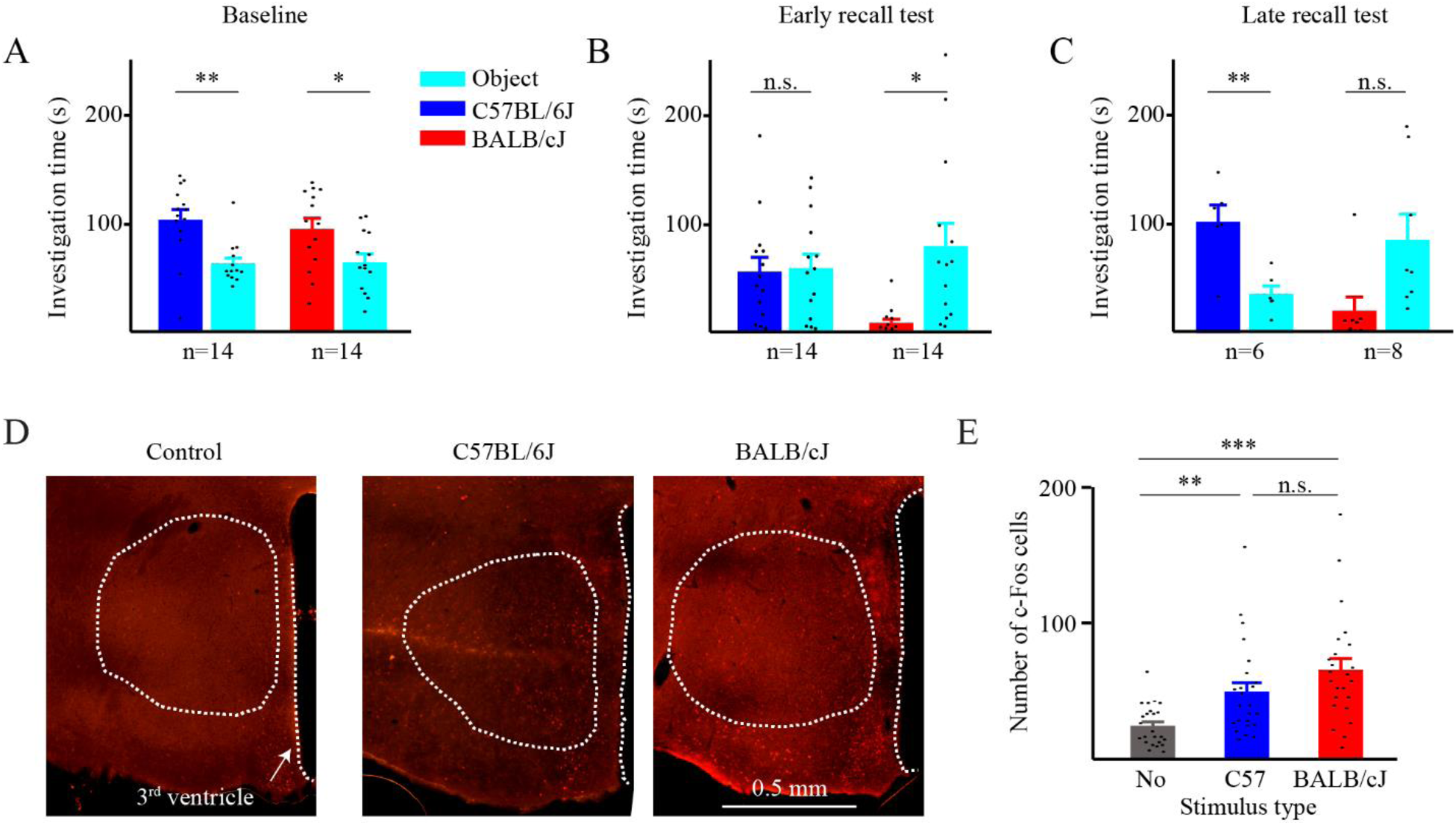
No difference in AHN c-Fos expression following affiliative vs. aversive social interaction. (A) Mean (±SEM) investigation time during Baseline SP tests conducted with either C57BL/6J or BALB/cJ social stimuli before SFC. (B) As in A, for Early recall tests. (C) As in A, for Late recall tests. (D) Representative images depicting c-Fos immunostaining in the AHN of animals after a Late recall test conducted using either no social stimulus (control, left), C57BL/6J social stimulus (middle), or BALB/cJ social stimulus (right). Note the similar number of stained cells in the latter two cases. (E) Mean (±SEM) number of c-Fos expressing cells after Late recall tests using either no social stimulus (control, gray), C57BL/6J social stimulus (middle, blue) or BALB/cJ social stimulus (right, red). Note that both social stimuli elicited a similar significant increase in c-Fos-positive cell number, as compared to the control. **p<0.01, ***p<0.001, paired t-test with Bonferroni correction for multiple comparisons following main effect in one-way ANOVA, or Dunn’s multiple comparisons following main effect in Kruskal Wallis test for non-parametric comparisons.

### Optogenetic stimulation of AHN neurons in a social context enhances approach behavior

To directly assess the role of the AHN in social approach or avoidance behaviors, we used optogenetic stimulation of AHN neurons in mice that were injected with a ChR2.0-expressing AAV viral vector and chronically implanted with optic fibers into the AHN (Fig. 6A-B). In slice whole-cell patch recordings, infected AHN neurons responded to 1-ms optogenetic stimulation with spiking activity that followed the stimulation frequency quite well between 5-20 Hz stimulation, but not so well above 20 Hz (Fig. 6C-D). Interestingly, AHN neurons were characterized with a prominent afterdepolarization component, which in many cases generated firing of spike doublets (Fig. 6C).

**Figure 6.**
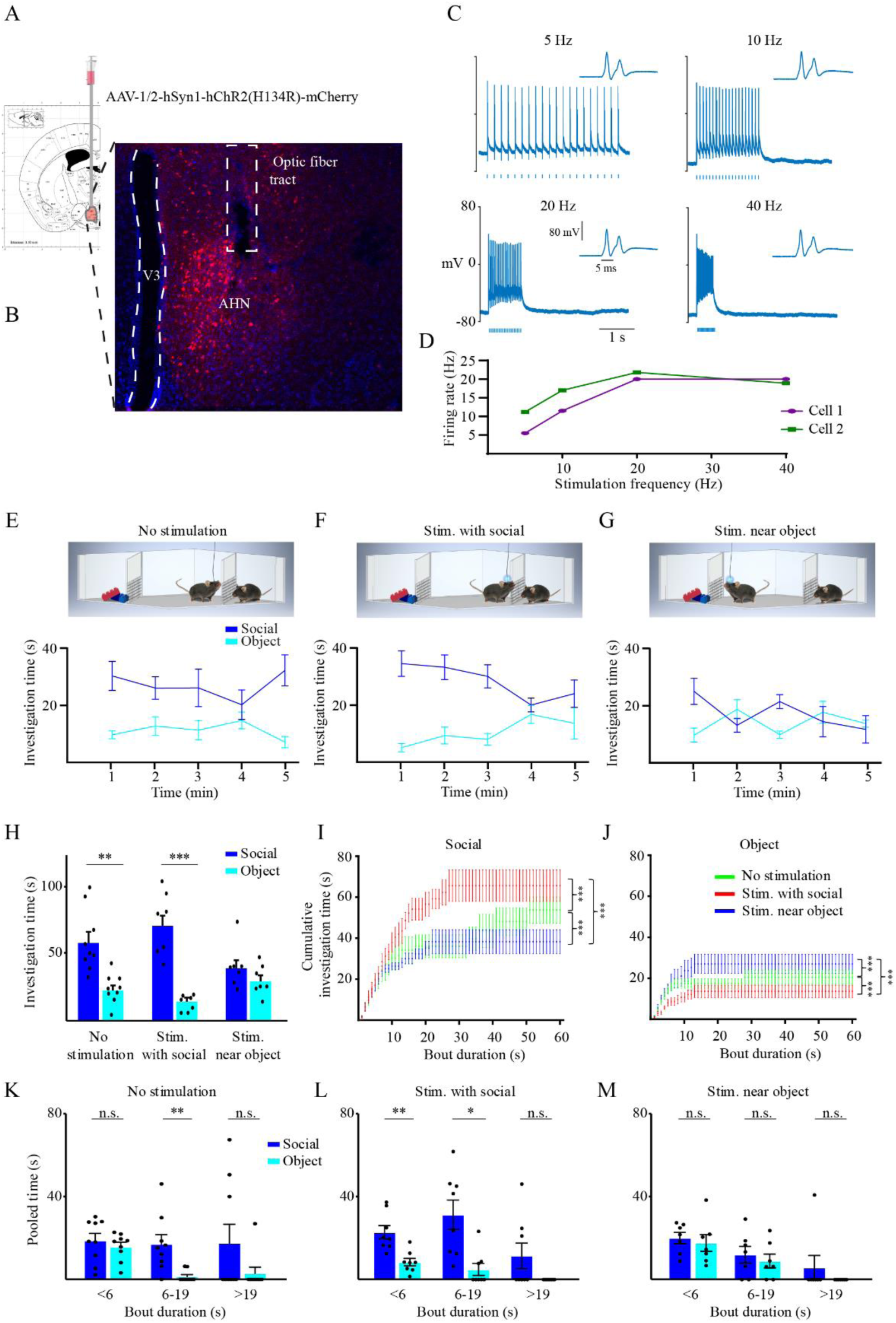
Optogenetic activation of AHN neurons enhances investigation of the stimulus associated with the activation. (A) A scheme of the viral injection to the AHN. (B) Representative image of AHN in a brain slice, depicting the infected neurons and optic fiber location. (C) Electrophysiological traces exemplifying whole-cell patch recording from a viral-infected AHN neuron in response to optogenetic stimulation at various rates (noted above the traces). The inset on the right of each trace shows the response to the first stimulus in higher time resolution, exposing a spike doublet. (D) The mean firing rate across the stimulation epoch, as a function of the stimulation frequency. Note the linear relationship between 5-20 Hz stimulation. (E) Mean (±SEM) investigation time across the SP session (1-min bin) for each stimulus, plotted for the no optogenetic stimulation (control) condition (schematically depicted above). Note the clear social preference exhibited by the subjects. (F) As in E, for optogenetic stimulation given during social investigation bouts. (G) As in E, when stimulation was given near the object (away for the social stimulus). Note the loss of social preference specifically in this condition. (H) Mean (±SEM) investigation time throughout the session for the three conditions shown in E-G. **p<0.01, ***p<0.001, paired t-test following main effect in ANOVA. (I) Cumulative histogram (mean±SEM) of social investigation time plotted as a function of bout duration, separately for the three stimulation conditions (color-coded). Note the statistically significant differences between the three curves. (J) As in I, for object investigation bouts. Note the inverse relationship between the curves as compared to I. (K) Mean investigation time pooled separately for short (<6 s), intermediate (6-19 s), and long (>19 s) investigation bouts, for each of the stimuli during control condition. Note the insignificant difference between social and object stimuli for short bouts. *p<0.05, paired t-test following main effect in ANOVA. (L) As in K, for stimulation during social investigation. Note the significant difference between social and object stimuli even for short bouts. *p<0.05, **p<0.01, paired t-test following main effect in ANOVA. (M) As in K, for stimulation near the object. Note that none of the bout duration showed any significant difference between the social and object stimuli. *p<0.05, **p<0.01, paired t-test following main effect in ANOVA. See also Fig. S5.

Four weeks after viral infection and implantation of an optic fiber, mouse subjects (n=9) were subjected to SP tests in which the animals were either unstimulated (Fig. 6E), stimulated at the beginning of social investigation bouts (Fig. 6F), or stimulated when being near the object stimulus (Fig. 6G). We did not restrict this latter condition to object investigation bouts only, since there are usually only few such events in each session. The three types of experiments were randomly conducted with the same animals, each on a separate day. Optogenetic stimulation was given on average 13 times in each type of stimulation conditions, with no significant difference between them (Independent samples t-test, t(13)=-0.308, p=0.763). When analyzing the investigation time dedicated to each of the stimuli across the various stimulation conditions, we found a significant interaction between stimulus and condition [two-way mixed model ANOVA-Stimulus: F(1,21)=38.41, p<0.001; Condition: F(2,21)=1.447; p=0.258; Stimulus x Condition: F(2,21)=5.579, p<0.05]. As expected, unstimulated animals exhibited a clear social preference throughout the test (Fig. 6E, H; Paired t-test: t(8)=3.585, p<0.01). Similar results were observed with animals that were stimulated during social investigation (Fig. 6F, H; Paired t-test: t(7)=5.755, p<0.001). In contrast, animals that were stimulated while being near the object stimulus, showed no social preference (Fig. 6G, H; Paired t-test: t(6)=1.293, p=0.243). Similar results were obtained when we analyzed data obtained only from animals that passed through all stimulation conditions (Fig. S4A-D, n=6). Thus, stimulating AHN neurons near the object stimulus prevented the mice from exhibiting their innate tendency to prefer social stimuli over object ones.

To further analyze these data, we plotted the cumulative investigation time as a function of investigation bout duration in all three conditions, separately for each stimulus. As apparent, there was a significant difference between the three conditions for both stimuli [Fig. 6I-J; *Social Stimulus:* Kolmogorov-Smirnov Z test: No stim. vs. Stim. with social: z=4.108, p<0.001; No stim. vs. Stim. away from social: z=2.556, p<0.001; Stim. with social vs. Stim. away from social: z=4.564, p<0.001; *Object stimulus:* Kolmogorov-Smirnov Z test: No stim. vs. Stim. with social: z=5.112, p<0.001; No stim. vs. Stim. away from social: z=4.747, p<0.001; Stim. with social vs. Stim. away from social: z=5.112, p<0.00]. Interestingly, optogenetic stimulation of AHN neurons near the social stimulus shifted the social investigation towards a higher number of short bouts, compared to both other conditions (Fig. 6I-J). We therefore summed the investigation time dedicated to each stimulus separately for short (< 6 s), intermediate (6-19 s), and long (> 19 s) bouts, as we have previously described (Netser et al., 2017). In accordance with our previous results, unstimulated animals did not show any difference between stimuli in short bouts, and their social preference was reflected only in investigation bouts which were longer than 6 s [Fig. 6K; two-way repeated measures ANOVA: Stimulus: F(1,8)=15.89, p<0.01; Bout duration: F(2,16)=1.046, p=0.374; Stimulus X Bout duration: F(2,16)=1.261, p=0.31; *post hoc* paired t-test: Short bouts: t(8)=1.167, p=0.277; Intermediate bouts-t(8)=3.458, p<0.01, Long bouts: t(8)=1.688, p=0.13]. In contrast, the social preference of animals stimulated near the social stimulus was reflected by both short and intermediate bouts [Fig. 6L; two-way repeated measures ANOVA: Stimulus: F(1,7)=26.995, p<0.001; Bout duration: F(2,14)=4.511, p<0.05; Stimulus X Bout duration: F(2,14)=1.268, p=0.312; *post hoc* paired t-test: Short bouts: t(7)=4.159, p<0.01; Intermediate bouts: t(7)=2.917, p<0.05, Long bouts: t(7)=1.865, p=0.104]. Animals stimulated near the object did not show any social preference [Fig. 6M; two-way repeated measures ANOVA: Stimulus: F(1,6)=1.637, p=0.248; Bout duration: F(2,12)=6.558, p<0.05; Stimulus X Bout duration: F(2,12)=0.122, p=0.886; *post hoc* paired t-test-Short bouts: t(6)=0.517, p=0.624; Intermediate bouts: t(6)=0.604, p=0.568, Long bouts-t(6)=1, p=0.356].

Thus, optogenetic stimulation of AHN neurons changed the subject’s investigation pattern in favor of the stimulus associated with the stimulation. Altogether, these results support a role for AHN neural activity in regulating approach behavior in a social context.

## Discussion

The AHN is a poorly defined and relatively unexplored brain area which is part of the hypothalamic defensive system (Canteras, 2002). Several previous studies that directly examined the effect of stimulating or inhibiting AHN neurons have yielded contradicting results. While some of them concluded that AHN stimulation cause escape response (Bang et al., 2022; Wang et al., 2015), others suggested the opposite (Anthony et al., 2014). Yet, none of these studies examined the role of the AHN in a social context. Interestingly, a recent study demonstrated that activation of vGAT-positive cells, a population consisting about 90% of AHN neurons, in a context of mechanical stimulation, elicits a robust biting attack by the stimulated mice (Xie et al., 2022). Such an attack may be considered the opposite response to escape, as it requires approaching the threat rather than avoiding it. Notably, the same study found that in a social context, AHN stimulation reduces aggression and enhances social investigation. These results motivated us to examine the role of AHN neurons in a social context.

Here we combined *in vivo* electrophysiology in behaving mice with c-Fos staining and optogenetic stimulation to probe the function of AHN neurons in social behavior. Specifically, we examined their role in regulating approach and avoidance behaviors during various social tasks. Overall, our results suggest that in social contexts, synchronized activity of AHN neurons is associated with approach behavior.

Using chronically-implanted tetrodes we measured, several distinct electrophysiological parameters along the social encounter: theta and gamma rhythmicity of LFP signals as well as the rate and theta rhythmicity of spiking activity. Our results suggest that the social context is reflected mainly by LFP theta rhythmicity, which was elevated during affiliative interaction and was reduced during aversive interaction. These results are in accordance with our previous results in rats (Tendler and Wagner, 2015), which demonstrated a social-context-induced elevation of theta rhythmicity in multiple limbic areas of the rat brain, including the medial amygdala (MeA) and LS, both of which project to the AHN. Yet, LFP theta and gamma power in the AHN almost did not change during investigation bouts towards the various stimuli, hence does not seem to be associated with specific behavioral events, but rather with the social context. These results differ from the results of similar recordings we have performed in the murine MeA, where we have recently reported that theta and gamma rhythmicity are also associated with social investigation bouts (John et al., 2022). Thus, our data suggest distinct roles for rhythmic neural activity in the MeA and AHN of adult male mice during social behavior.

In contrast to LFP theta and gamma rhythmicity, we found spiking activity in the AHN to be generally reduced in a social context. Such general reduction in spiking activity may support enhanced signal-to-noise ratio (SNR) while the AHN networks process information acquired during the investigation bout, a phenomenon well-known from other brain areas (Barta and Kostal, 2019; Ferguson and Cardin, 2020). Nevertheless, we found that spiking activity in the AHN is specifically modified during investigation behavior, in a stimulus-dependent manner. While during object investigation, the spiking response rapidly declined below baseline, during social investigation, it was kept above baseline for several seconds. This significant difference between the two types of stimuli may be related to the fact that investigation bouts towards social stimuli tend to be longer than towards objects (Netser et al., 2017). Nevertheless, since the subject animals showed a robust preference for investigating the social stimulus as compared to the object, the higher spiking activity of AHN neurons during social investigation bouts strongly suggests that in a social context, AHN neuronal activity is associated with approach behavior rather than avoidance. These results are in agreement with the results of both Anthony et al. (2014) and of Xie et al. (2022). Moreover, they do not contradict the multiple studies conducted with various rodent species, which implicated the AHN in aggressive behavior (Gobrogge et al., 2007; Haller et al., 2006; Ricci et al., 2009; Tachikawa et al., 2013), as such behavior also requires approach towards social stimuli. Thus, AHN neuronal activity may enhance approach towards social stimuli, regardless of whether in affiliative or aggressive manner.

Employing a novel behavioral paradigm of stimulus-specific SFC, we examined the responses of AHN neurons to attractive and aversive social stimuli. Our c-Fos analysis indicates a similar number of AHN neurons activated during encounters with both types of social stimuli. One possibility is that distinct neuronal ensembles are activated by the opposite types of social stimuli, one of which is associated with approach behavior while the other triggers avoidance. It should be noted that such an arrangement was recently demonstrated in the PAG (Reis et al., 2021), one of the main targets of the AHN (Hahn and Swanson, 2015; Semenenko and Lumb, 1999). Alternatively, the same neurons may be activated by both stimuli, but in a different manner. Notably, we found that in social contexts, AHN spiking activity exhibited enhanced theta rhythmicity and was synchronized with the enhanced LFP theta rhythmicity. Thus, both attractive and aversive social stimuli may trigger similar firing rates of AHN neurons, hence a similar level of c-Fos expression. Yet, in the affiliative social context, the enhanced LFP theta rhythmicity may impose (or reflect) more coherent spiking activity in the AHN, which would drive the animals towards social approach. In contrast, non-coherent AHN activity driven by aversive stimuli may fail to induce approach behavior or even elicit avoidance.

In accordance with this suggestion, the coherent activation of AHN neurons by our optogenetic stimulation drove the animals to interact with the stimulus associated with optogenetic activation, regardless of its nature (social or object). It should be noted that in previous studies, direct optogenetic stimulation of AHN neurons caused distinct responses, in a context dependent manner. These included escape in an empty arena, biting attack following a mechanical stimulation and enhanced social investigation in a social context. In light of all these studies, our results further suggest that the behavioral effect of AHN activation is highly context-dependent. How exactly does the type of context affect the consequences of AHN activity should be explored in future studies. Yet, we suggest that one mechanism which may be involved is the type and level of theta rhythmicity and the synchronized activity it may impose on various brain regions. It should be noted that in our previous study where we used multi-site brain recording from behaving rats we found that social and fear stimuli induce distinct types of theta rhythmicity and different patterns of coherence in some regions of the SBN (Tendler and Wagner, 2015). Such modulation of theta rhythmicity and its coherence may therefore encode various emotional contexts and guide distinct outcomes of activity in the relevant brain regions.

## Declarations

## Acknowledgments

We thank Boris Shklyar, Head of Bioimaging Unit and Alex Bizer, the experimental systems engineer of the Faculty of Natural Sciences of the University of Haifa, for their help. We also thank Dr. Yair Shemesh from the Weizmann Institute, Rehovot, Israel, for critical reading of the manuscript draft.

## Author contributions

Conceptualization (SN, SW), Methodology (SN), Software (SN), Validation (SN, SW), Formal analysis (RJ, SN, SP, WD), Investigation (BKP, RJ, SN, SP, WD), Resources (EB), Writing – original draft ((RJ, SN, SW), Writing – review & editing (EB, SN, SW), Visualization (RJ, SN), Supervision (SW), Project administration (SN), Funding acquisition (SW).

## Availability of Data and Materials

The datasets used and/or analyzed during the current study as well as the detailed statistical analyses are deposited in Mendeley Data and available using the following reference - Netser, Shai (2022), “Data and statistical summary for the paper - Neural activity and theta rhythmicity in the anterior hypothalamic nucleus are involved in regulating social investigation”, Mendeley Data, V1, doi: 10.17632/k6htkkfz42.1

## Methods

### Animals

Mouse subjects were naïve C57BL/6J adult male mice (8-18 weeks old), commercially obtained (Envigo, Israel). Social stimuli were in-house grown C57BL/6J juvenile male mice (3-6 weeks old) in the SP and free interaction tests, or adult male C57BL/6J, BALB/cJ or CD-1 (ICR) mice in the case of SFC paradigm. All animals were kept in groups of 2-5 per cage at the animal facility of the University of Haifa under veterinary supervision, in a 12 h light/12 h dark cycle (lights on at 9 PM), with *ad libitum* access to food (standard chow diet, Envigo RMS, Israel) and water. In experiments involving electrodes or optic fiber implantation, mice were kept in isolation for about 7 days following surgery.

### Experiments

Behavioral experiments took place during the dark phase of the animals, under dim red light. All experiments were performed according to the National Institutes of Health guide for the care and use of laboratory animals and approved by the Institutional Animal Care and Use Committee (IACUC) of the University of Haifa.

### Experimental setups

#### Social preference setup

The experimental setup was as previously described (Netser et al., 2019). Briefly, the setup consisted of a white Plexiglas arena (37 X 22 X 35 cm) placed in the middle of an acoustic chamber (60 X 65 X 80 cm). Two Plexiglas triangular chambers (12 cm isosceles, 35 cm height) were placed in two randomly selected opposite corners of the arena, in which an animal or object (plastic toy) stimulus could be placed. A metal mesh (12 X 6 cm, 1 X 1 cm holes) placed at the bottom of the triangular chamber allowed direct interaction with the stimulus through the mesh. A high-quality monochromatic camera (Flea3 USB3, Point Grey), equipped with a wide-angle lens, was placed at the top of the acoustic chamber and connected to a computer, enabling a clear view and recording (∼30 frames/s) of the subject’s behavior using a commercial software (FlyCapture2, Point Grey).

#### Free interaction setup

Following the SP test, the chambers separating between the subjects and the social stimuli were removed from the arena, and a free interaction session was conducted in the same arena.

#### Social fear conditioning setup

The SFC setup was a custom-made white Plexiglas arena similar in size to the experimental setup for the social preference test (37 X 22 X 35 cm) but with a metal grid floor (H10-11M, Coulbourn Instruments) connected to an electrical shock-delivering unit (precision regulated animal shocker H13-14, Coulbourn Instruments). The unit was modified to deliver a single pulse of 0.3-0.4 mA for 750 ms when manually triggered.

### Behavioral paradigms

#### Social preference/free interaction paradigm used for electrophysiology

After connecting the subject to the recording system, it was left for a 15-min habituation to an arena containing empty chambers. Throughout this time, social stimuli were placed in similar chambers near the acoustic chamber for acclimation. The recording session started with an additional 5 min of baseline (pre-encounter) period with the empty chambers. Thereafter, the empty chambers were replaced with chambers containing the social and object (plastic toy, ∼5x5 cm) stimuli, and the SP test was conducted for 5 min. Following the SP test the stimuli-containing chambers were replaced again with the empty ones for an additional 5 min of recordings (post-encounter, see Fig 1A). Then, the empty chambers were removed from the arena, and the subject was left alone for 20 min. Next, the recording started again for an additional 5 min of baseline (pre-encounter), followed by a 5-min test of free interaction with a novel social stimulus taken directly from its home cage and an additional 5 min of post-encounter period after removal of the social stimulus back to its home-cage. Each subject (n=17) animal was tested 2-6 times, with at least 6 h separating between distinct sessions with the same subject.

#### Social preference used for in vivo optogenetic stimulation

After connecting the subject (n=9) to the stimulation apparatus, it was left for a 15-min habituation to an arena containing empty chambers. The interaction was recorded for 5 min, after replacement of the empty chambers with the social and object stimuli chambers. Each subject was tested three times – unstimulated (Fig. 6E), stimulated at the beginning of social investigation bouts (Fig. 6F), or stimulated when being near the object stimulus (Fig. 6G). The three types of experiments were randomly conducted with the same animals, each on a separate day.

#### Social Fear Conditioning paradigm

The SFC paradigm consisted of a 15-min habituation of the subjects to an arena containing two empty chambers, followed by two consecutive SP tests with social stimuli of two distinct strains (C57BL/6J and either ICR or BALB/cJ), separated by a 15-min interval. Thereafter, the subject was transferred to the SFC arena for a 15-min habituation, followed by 5 min of the SFC procedure, in which the subject received a mild electrical foot shock (0.3-0.4 mA, 750 ms) each time it tried to interact with the stimulus chamber (ICR or BALB/cJ strains). Five minutes after conditioning, the subject was returned to the experimental arena for a 15-min habituation and two more consecutive SP tests, as before conditioning (Early recall tests). The animals were then placed back in their home-cages and a day later were returned to the experimental arena for a 15-min habituation, and two more consecutive SP tests were performed (Late recall tests). In sessions with electrophysiological recordings (n=7), the animal’s behavior and brain activity were recorded for 5 min of pre-encounter (with empty chambers), 5 min of encounter (with the social stimulus), and 5 min after the encounter (with empty chambers). In sessions in which the animals were sacrificed for c-Fos immunostaining (n=14, out of which four from each group were randomly taken for analysis), the Late recall test was done only once, with one of the social stimuli (C57BL/6J or BALB/cJ).

### Tracking software and behavioral analyses

All recorded video clips were analyzed using TrackRodent (https://github.com/shainetser/TrackRodent) as previously described (Netser et al., 2019). For analysis of SP sessions, we used the *BlackMouseWiredBodyBased* algorithm and for free interaction sessions we employed the *BlackMice_TwoMiceFreeInteraction* algorithm as previously described (John et al., 2022).

### Electrode implantation surgery

Mice were anesthetized and analgized with an intraperitoneal injection of a mixture of ketamine, Domitor, and the painkiller Norocarp (0.13 mg/gr, 0.01 mg/gr, and 0.005 mg/gr, respectively). Anesthesia level was monitored by testing toe pinch reflexes and additional doses (30% of the initial dose) were administered if necessary. The body temperature of the animals was kept constant at approximately 37°C, using a closed-loop custom-made temperature controller connected to a temperature probe and a heating pad placed under the animal. Anesthetized animals were fixed in a stereotaxic apparatus (Kopf Inst.), with the head flat. The skin was then gently removed, and holes were drilled in the skull for implanting a custom-made tetrode probe, fixing supporting screws and placing reference and ground silver wires. The custom-made probe consisted of 4 tetrodes (16 Ni-chrome wires, polyimide insulated, 12µm diameter core bare at the tip, AM systems) glued to an optic fiber (320µm with coating, Thorlabs) and fixed to a custom-made 3D plastic-printed scaffold (See Fig. S1A). The tip of each wire was inserted to a 9*2 Mill-Max female connector (Interconnect Machined Pin Socket, #853-43-100-10-001000, Mill-Max) stripped of pins, which were then inserted back to the socket by manually pressing them into it. The assembly was then glued on a miniature custom-made drive (Vandecasteele et al., 2012) which enabled driving the probe deeper into the brain between sessions if no spikes were detected. The wires were then plated with Platinum Black Plating Solution (Neurolynx) to lower the impedance down to a range of 200 KΩ to 350 kΩ (measured using LCR meter, Tonghui). The additional two pins were soldered to silver wires (100 µm silver wire, AM systems) and served for reference and ground. Before implantation, electrodes were coated with a dye, DiI (1,1’-Dioctadecyl-3,3,3’,3’-tetramethylindocarbocyanine perchlorate, dissolved in 70% ethyl-alcohol; Invitrogen), for fluorescent marking aimed to track their position post-mortem. Tetrodes were implanted above the AHN (A/P = − 0.85 mm, L/M = − 0.2 mm, D/V = − 4.75 mm) and were driven into the AHN before the recording sessions. The drive, wires, and screws were fixed by dental cement and the implant was covered with copper foil plates, which were connected to the ground wire as well (Vandecasteele et al., 2012). After the surgery, Antisedan was given subcutaneously to wake the mice from the anesthesia. The mice were then placed in a warm cage (37°C) overnight. The animals were injected with Norocarp and Baytril (5%, 0.03 ml/10gr) to relieve pain and prevent infections for three days following the surgery. Behavioral testing and electrophysiological recordings were conducted at least five days post-surgery.

### Electrophysiological recordings

As previously described (John et al., 2022).

### Analysis of LFP signals

All signals were analyzed using a custom-made MATLAB program. First, the signals were down-sampled to 5 kHz and low-pass filtered up to 0.3 kHz using a Butterworth filter. The power over time for the different frequencies was constructed by the “spectrogram” function in MATLAB, using a 2 s long discrete prolate spheroidal sequences (DPSS) window with 50% overlap, at 0.5 Hz increments and 0.5 s time bins. The power for each frequency band (Theta: 4-12 Hz and Gamma: 30-80 Hz) was averaged for each recorded channel. The channel chosen for further analysis was the one with the highest and most stable signal to noise ratio (SNR) in the specific frequency (Theta or Gamma) during the post-encounter period (mean divided by standard deviation of the signal).

For evaluation of the change in activity across the entire recording session, the signal was Z-scored by subtraction of the mean power and division by the standard deviation of the pre-encounter period. For presentation, the average Z-score was calculated in 10 s bins.

Synchronization between the LFP signal and investigation bouts towards social or object stimuli was assessed by calculating the power within a time window of 5 s before and 6 s after the beginning of each bout (using 1 s bins) and then averaging the responses to all bouts for each session. Thereafter, the mean power was normalized separately for each session using Z-score analysis, with the 5 s before the bouts serving as a baseline.

### Analysis of spiking activity

All signals were analyzed using a custom-made MATLAB program. First, the signals were band-pass filtered from 0.3 kHz to 5 kHz using a Butterworth filter. Then, the time of spikes in each channel was detected using a threshold of 3.5 standard deviation (calculated for each minute of the recording separately). We then sorted the spikes into single and multi-units based on the data from each tetrode using PCA analysis based on the spike shape and amplitude using the software package written by Alexei A Koulakov (Shusterman et al., 2011). Since many of the spikes could not be sorted well into single units, we considered all units as multi-unit activity.

The change in multi-unit activity across the entire recording session and its synchronization with the investigation bouts were analyzed in a similar manner to the LFP analysis. For presentation, the average Z-score was calculated in 30 s bins.

### Spike-triggered LFP (stLFP) analysis

We calculated stLFP by averaging short segments of the LFP signal aligned with each spike. First, raw LFP signals were band-pass filtered at the Theta range and normalized by subtracting the mean and dividing by the standard deviation of the entire signal recorded. Next, short segments of 200 ms, centered around each spike occurring during the encounter, were averaged for each unit separately. In addition, we averaged a similar number of segments around random time points within the encounter time (shuffled-LFP). For statistical comparison between the real and shuffled data, we summed the stLFP and shuffled-LFP average at the time window of -10 to +10 ms around the spike time. It should be noted that more than 1500 spikes were recorded from each multi-unit, therefore no multi-unit was excluded.

### *In vivo* optogenetics

#### Optic fiber

To enable *in vivo* delivery of light, optic fibers were prepared in-house by inserting 200 mm optic fibers with 0.39 N.A. into a 10.5 mm long ceramic ferrule with outer diameter of 2.5 mm (Thorlabs), gluing them in place, and polishing the ferrule tip with 30, 6, and 3 µm sandpapers (Thorlabs). The optic fiber tip was then cut to an appropriate length that is sufficient to reach the AHN. Light power at the tips of the fibers was measured for adequate power output (∼10 mW) using a power meter (Thorlabs) prior to implantation. This parameter was later used to determine the driving voltage needed for achieving identical stimulation power in all cases.

#### Surgery for virus injection and optic fiber implantation

Subject mice were first anesthetized as described above. The mouse’s head was then shaved and fixed in a stereotaxic apparatus (Kopf Inst.). The scalp was cut to reveal the skull, and measurements were then taken to ensure alignment of the skull. The AHN was targeted according to the coordinates described above for *in vivo* electrophysiology. Once the coordinates were marked, a hole (unilateral, right hemisphere) was drilled, and a glass capillary filled with the virus (ssAAV-1/2-hSyn1-hChR2(H134R)_mCherry-WPRE-hGHp(A), Zurich VVF, Switzerland) was slowly lowered into the target region and left in place for 5 min prior to injection and 10 min following injection to prevent retraction of the virus. A total of 300 nl of the virus was then delivered by manual application of pressure using a 50 ml syringe connected to the glass capillary (BRAND, disposable BLAUBRAND® micropipettes, intraMark, 5*μl*). Following the viral injection, an optic fiber was inserted into the region of interest with the same coordinates as the viral injection and placed 100 *μ*m above the injection depth. After the surgery, Antisedan (0.1ml/10gr bodyweight) was given subcutaneously to wake the mice from the anesthesia. The mice were then placed in a warm cage (37°C) overnight. The animals were injected with Norocarp and Baytril (5%, 0.03 ml/10gr) to relieve pain and prevent infections for three days following the surgery. Behavioral testing was conducted three weeks after the viral injection.

#### Optogenetic stimulation

On the day of testing, mice were lightly anesthetized with isoflurane and connected to the laser (473 nm) for light delivery. Light power was set at 5 mW, which is sufficient to yield 1 mW/mm^2^ - the minimum amount of light reported to be capable of efficient ChR2 activation (Knobloch et al., 2012), positioned no farther than 0.25 mm above the AHN. This should lead to a specific stimulation of the AHN, as previous measurements with blue light stimulations in rodent brains have shown that the blue light of the laser does not penetrate the tissue further than 500 μm (Yizhar et al., 2011). Each animal underwent three sessions of SP on separate days as described above. The optical stimulation in all cases was a 20 Hz train of 20 ms stimuli given for 1 s. These parameters were chosen in accordance with the study of Anthony et al. (2014). The optical stimulation was manually delivered via a Master8 Channel Programmable Pulse Stimulator (A.M.P.I).

#### Post-mortem histological location analysis of electrodes and optic-fibers

Animals were perfused with phosphate buffer saline (PBS) and then fixed using 4% paraformaldehyde (PFA, Sigma) solution. The brains were harvested and placed in PFA (4%) for 48 hours, followed by sectioning of 50-µm slices on the horizontal axis using a VT1000s sliding vibratome (Leica.) Sections were stained with DAPI and examined under a wide-field fluorescence microscope (Nikon Ti-eclipse) for verifying the placement of the electrodes’ marks (DiI fluorescence) or the optic fiber within the AHN (Fig. S1C).

### *In vitro* electrophysiology

Two C57BL/6J adult male mice were injected with the viral vector as described above and kept in their home-cages for 4-8 weeks until their brains were harvested by decapitation following anesthesia with isoflurane. Fresh coronal brain slices (300 µm thick) were obtained in ice-cold oxygenated (95% O_2_-5% CO_2_) normal saline Ringer solution (in mM: 124 NaCl, 3 KCl, 2 MgSO_4_, 1.25 NaH_2_PO_4_, 26 NaHCO_3_, 2 CaCl_2_, and 10 glucose). AHN-containing slices were incubated for at least 1 h in room temperature. Then, the slices were transferred to a recording chamber perfused with Ringer solution maintained at 30°C, located beneath an infrared DIC microscope (BX51WI Olympus; Tokyo, Japan), with 10X or 40X water immersion objectives. Neurons expressing mCherry were identified for current-clamp recordings by their red fluorescence using excitation λ = 545-580 nm, emission λ ≥ 610 nm (fluorescence filter from Olympus, U-MWIY2).

Whole-cell current-clamp recordings were performed to measure the action potential firing rate using an Axopatch 200B amplifier and a 1550B digitizer (Molecular Devices, Sunnyvale, CA). Borosilicate glass pipettes (3–5 MΩ) were used with a 2-step electrode puller (Narishige PC-10) to make recording electrodes, which were then filled with internal solution (130 mm K-gluconate, 5 mm KCl, 10 mm HEPES, 2.5 mm MgCl_2_, 0.6 mm EGTA, 4 mm Mg-ATP, 0.4 mm Na_3_GTP, and 10 mm phosphocreatine: osmolarity 290 mOsm, pH 7.3). Data were obtained utilizing pClamp9 (Molecular Devices), acquired at 2 KHz and digitized at 5 KHz. Once the neuron was electrically accessible, the voltage data were obtained after the pipette capacitance and series resistance were compensated for manually. Before the recordings started, the blue-light pulse (excitation λ = 450-480 nm, ultra-high power led, Prizmatix) intensity was adjusted between 3 to 5 mW/mm^2^ until a single action potential was observed. Once optimized, the same light intensity was maintained throughout the recording of that cell. In order to validate the functionality of the virus *ex-vivo*, trains of 20 pulses (pulse duration 1 ms) were delivered at 5 Hz, 10 Hz, 20 Hz, and 40 Hz with a 30-s inter-train interval. Additionally, a train of 20-ms long pulses was delivered at 20 Hz. The light pulse train was delivered both sequentially (each trial containing all the frequencies) and each frequency with repeated trials.

### **c-** Fos expression

Ninety minutes following the Late recall test, mice were transcardially perfused with 100-200 ml of 0.1M Phosphate Buffered Saline (PBS) (pH=7.4), followed by ∼100 ml 4% Paraformaldehyde (PFA) in PBS. The brains were removed and kept in 4% PFA until sectioning. Coronal brain slices of 50 µm thickness were obtained as described above.

Free-floating brain sections were rinsed (3 x 10 min) in PBS and incubated in blocking solution [PBS+5% normal goat serum (Biological Industries) and 0.3% Triton X-100 (Fluka analytical)] for 2 hours at room temperature. Then, the sections were incubated with primary antibody against c-Fos [Anti-rabbit (Cell signaling) 1:500] at 4°c in a shaking incubator overnight. The sections were then rinsed again in PBS (3 x 10 min) and treated with secondary antibody [Alexa fluor 594-conjugatedanti-rabbit IgG antibody (Abcam), 1:500] for 2 hours at 4°c. After washing in PBS (3 x 10 min) at room temperature, sections were mounted on clean glass slides, air-dried, and cover slipped with DAPI + mountant (GBI labs).

### Microscopy

Fluorescence images were acquired through a 20X objective lens and a Zyla camera or a 10X objective and a Nikon camera, attached to the Nikon Ti2 fluorescent microscope. The images were acquired using a blue filter for DAPI staining, and TRITC for the c-Fos expressing cells. The image files were visualized using NIS Elements viewer software (Nikon) to manually count c-Fos-stained cells. Number of cells was an average over six randomly selected AHN slices per animal.

### Statistical analysis

All statistical tests were performed using SPSS v23.0 (IBM). The Kolmogorov–Smirnov test was used for verifying the normal distribution of the dependent variables. A two-tailed paired t-test was used to compare between different conditions or stimuli for the same group, and a two-tailed independent t-test was used to compare a single parameter between distinct groups. When the normal distribution assumption was violated, the non-parametric Wilcoxon test was used to compare between different conditions or stimuli for the same group, and Mann-Whitney for comparing between different conditions or stimuli for different groups. For comparison between multiple groups and parameters, a mixed analysis of variance (ANOVA) model was applied to the data. This model contains one random effect (ID), one within effect, one between effect, and the interaction between them. For comparison within a group using multiple parameters, a two-way repeated-measures ANOVA model was applied to data. This model contains one random effect (ID), two within effects, one between effect, and the interactions between them. All ANOVA tests were followed, if main effects or interaction found, by *post hoc* Student’s t-test with Bonferroni correction. For comparing between multiple groups when the normal distribution assumption was violated, Kruskal-Wallis test was used, followed by Dunn’s multiple comparisons for *post hoc* analysis. For testing main effects within a group for multiple parameters when the normal distribution assumption was violated, Friedman’s test was conducted. Significance was set at 0.05 and was adjusted when multiple comparisons were used.

## Supplementary Figures

**Figure S1.**
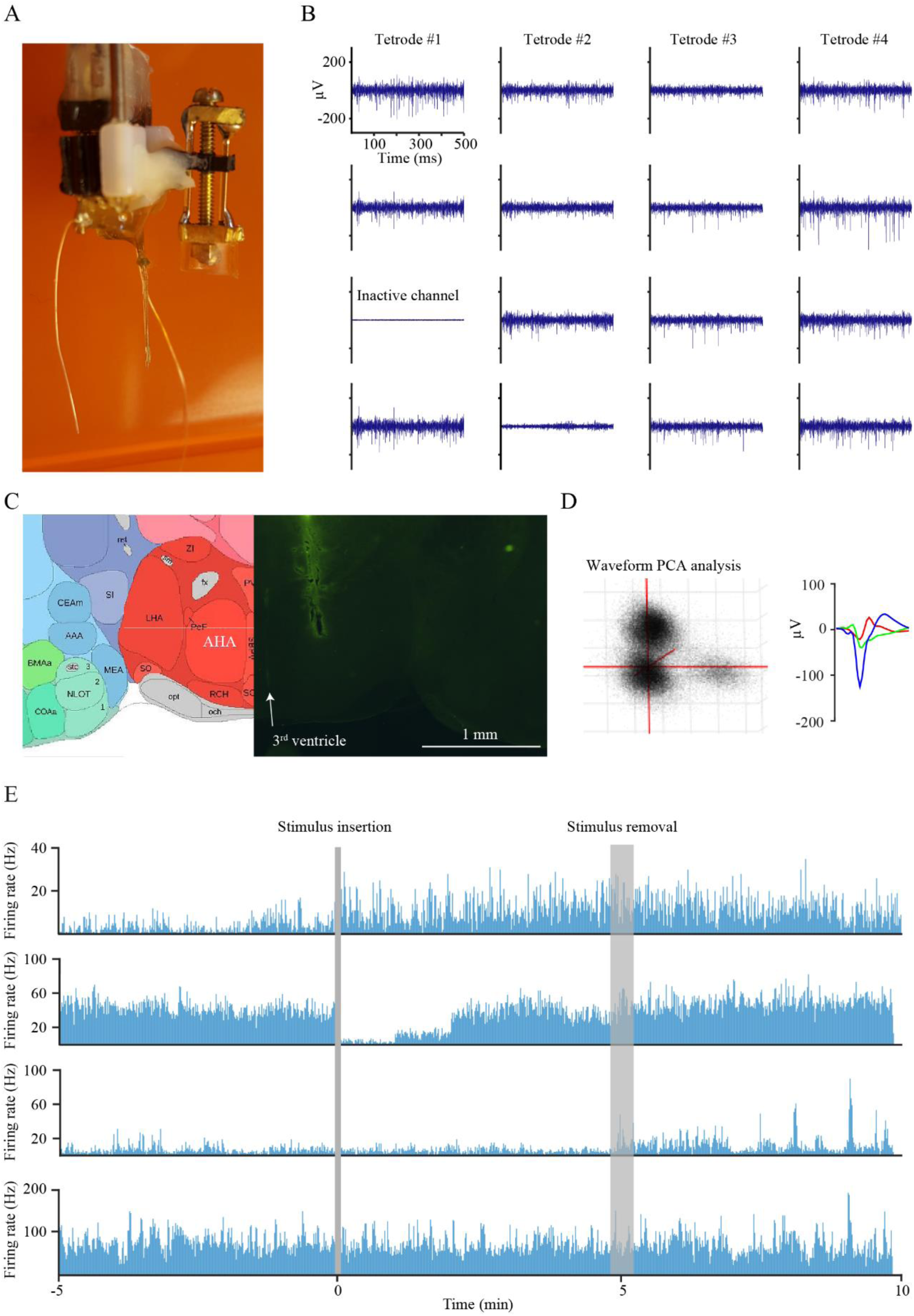
Recording AHN spiking activity. (A) A picture of the custom-made 4 x 4 tetrodes probe. (B) Representative 0.5-s traces from all 16 probes of one probe implanted into the AHN. (C) Implantation site of the tetrode probe: brain atlas (left) and representative image (right).Left: PCA analysis of a single recording session, showing the clusters created by three distinct single-units. Right: superimposed waveforms of the three units. (D) Firing rate of four distinct multi-units along the time course of the SP session. Gray stripes represent the time of stimulus insertion and removal.

**Figure S2.**
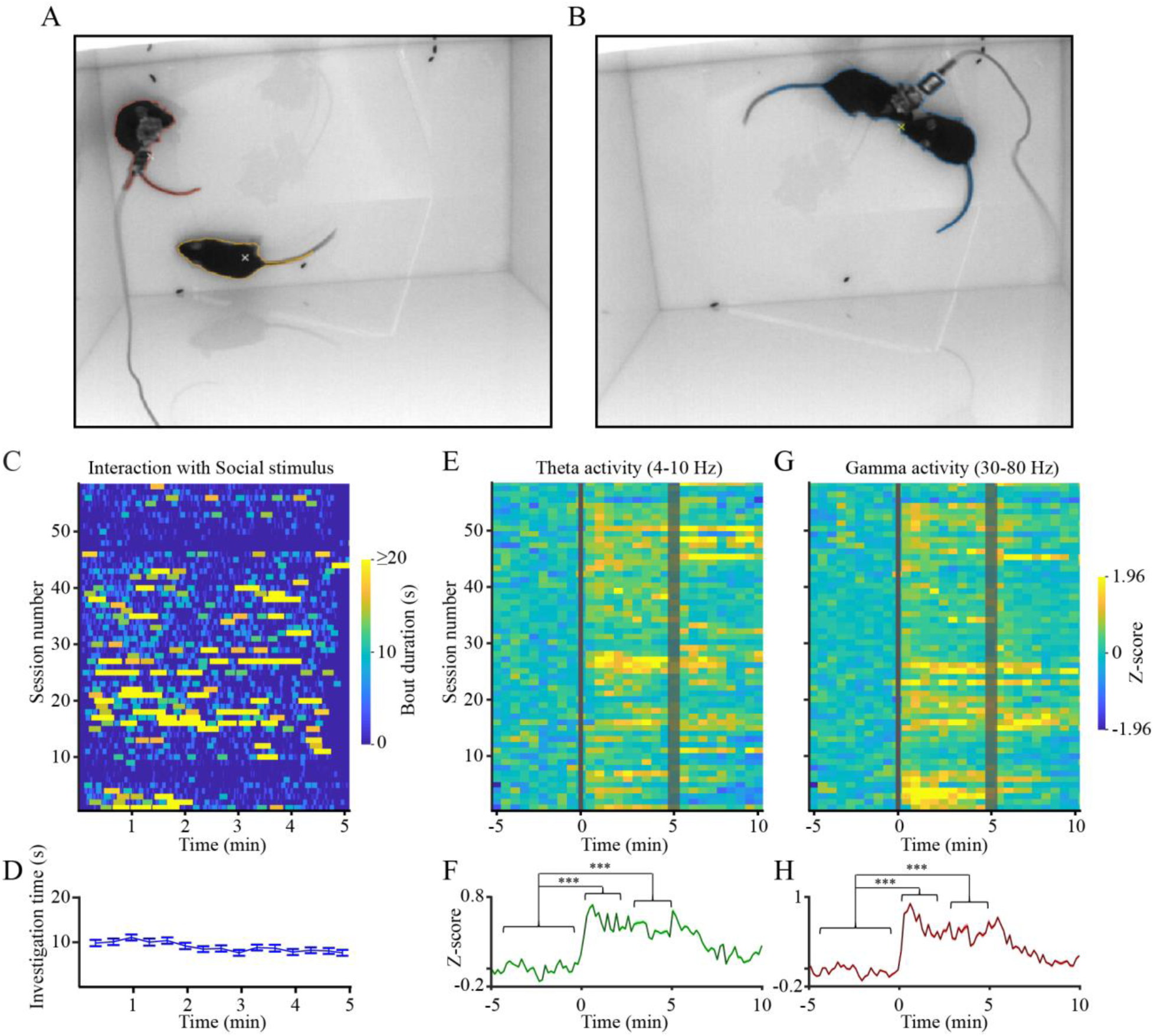
AHN theta and gamma rhythms are enhanced during free social interactions. (A) A picture of the experimental arena of the free social interaction paradigm with the two animals (subject + stimulus), while they are separated. (B) As in A, for the two animals while they are in contact. (C) Behavioral raster-plots of interaction (contact) bouts between the two animals along 56 sessions of free social interactions. (D) Mean interaction time (±SEM, 20-s bins) during the same sessions shown in C, along the time course of the free interaction session. (E) Heat-map of Z-score analysis of theta power of LFP signals recorded during the same sessions shown in C-D. Gray stripes represent the time of insertion (thin stripe) and removal (thick stripe) of the stimuli from the chambers. (F) Mean (±SEM) Z-score analysis of theta power of LFP signals recorded during the same sessions shown in D-E. (G) As in E, for gamma power. (H) As in F, for Z-score of gamma power. ***p<0.001, paired t-test with Bonferroni correction for multiple comparisons following main effect of time in ANOVA

**Figure S3.**
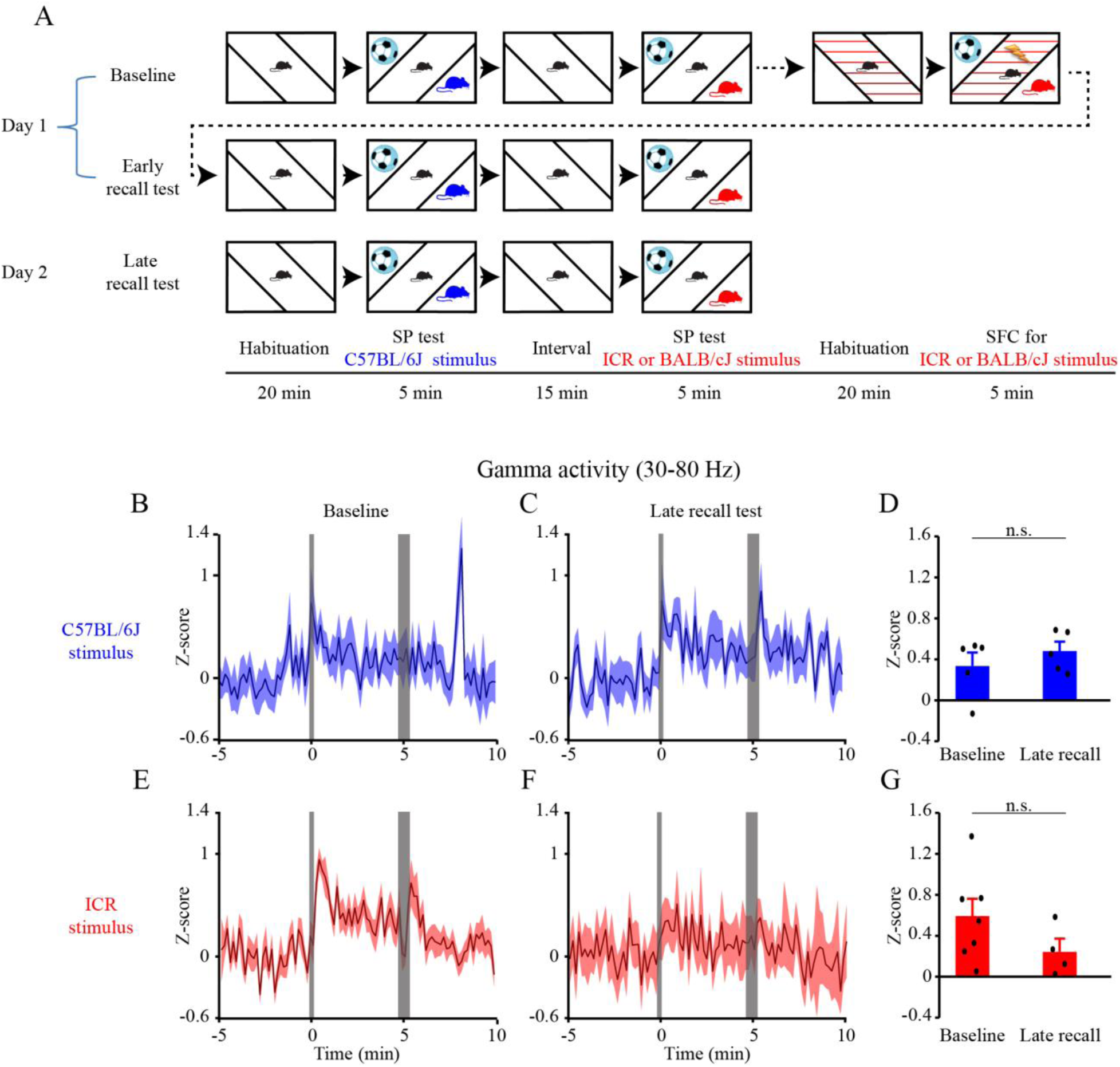
Gamma power of LFP signals in the AHN do not differ between affiliative and aversive social interactions. (A) A schematic depiction of the social fear conditioning (SFC) paradigm. Note that C57BL/6J stimuli are colored in blue while ICR stimuli are colored in red. (B) Mean (±SEM) Z-score analysis of gamma power along the time course of the Baseline SP test using C57BL/6J mice. (C) As in B, for the late recall test using the same stimulus. (D) Mean Z-score analysis of the change in gamma power during the Encounter stage of the SP test using C57BL/6J stimuli, calculated separately for Baseline (left) and Late recall (right) experiments. (E) As in B, for ICR stimuli. (F) As in C, for ICR stimuli. (G) As in D, for ICR stimuli.

**Figure S4.**
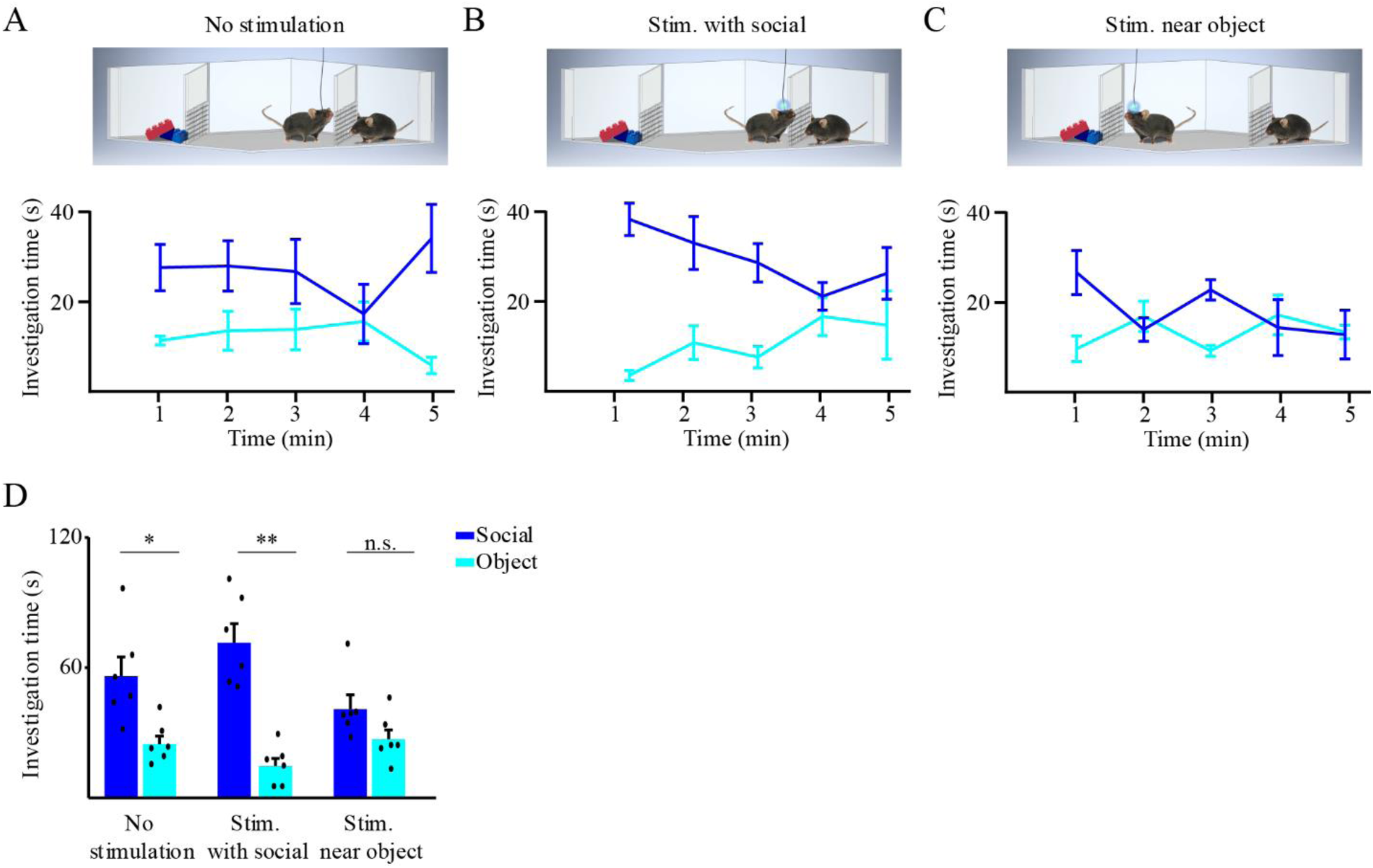
Optogenetic activation of AHN neurons enhances investigation of Optogenetic activation-associated stimulus, even when only the animals that passed all conditions are considered. (A) Mean (±SEM) investigation time across the SP session (1-min bin) for each stimulus, plotted for the no optogenetic stimulation (control) condition (schematically depicted above). Schematic depiction of the condition is presented above. (B) As in A, for optogenetic stimulation given during social investigation bouts. (C) As in A, when stimulation was given when the subject was near the object (away for the social stimulus). Note the loss of social preference specifically in this condition. (D) Mean (±SEM) investigation time throughout the session for the three conditions shown in A-C. *p<0.05, **p<0.01, paired t-test following main effect in ANOVA.

